# Dynamic and Heterogeneous Distribution of *Trichomonasvirus* Species in *Trichomonas vaginalis*

**DOI:** 10.1101/2025.03.31.646274

**Authors:** Hong-Wei Luo, Seow-Chin Ong, Jhen-Wei Syu, Chih-Yu Tsai, Po-Jung Huang, Chi-Ching Lee, Yuan-Ming Yeh, Rose Lin, Cheng-Hsun Chiu, Petrus Tang

## Abstract

*Trichomonasvirus* (TVV) is a double-stranded RNA virus from the *Pseudototiviridae* family that exclusively infects *Trichomonas vaginalis*, the protozoan responsible for the most common non-viral sexually transmitted infection. Within the *Trichomonasvirus* genus, five distinct viral species have been identified, and intriguingly, a single *T. vaginalis* isolate can simultaneously harbor multiple TVV species. Previous studies have explored the impact of TVV infection on the physiology and virulence of *T. vaginalis*, yet findings remain inconsistent and often contradictory. We propose that these discrepancies stem from the heterogeneous intracellular distribution of TVV within individual cells of the same *T. vaginalis* isolate. To investigate this, we employed ultra-deep single-cell RNA sequencing and immunofluorescence assays to examine multiple *T. vaginalis* isolates. Our results revealed striking variability in the proportion of infected cells carrying the same viral species, even within a single isolate. The present study is the first to demonstrate the dynamic and non-uniform intracellular distribution of TVV at the single-cell level. These findings challenge previous assumptions and provide a new perspective on the intricate virus-host interactions between TVV and *T. vaginalis*, offering new avenues for further research into their complex relationship.

**Author Summary:** *Trichomonas vaginalis* is a parasitic protozoan responsible for the most prevalent non-viral sexually transmitted infection worldwide. In 1985, researchers discovered a double-stranded RNA (dsRNA) virus within T. vaginalis, which was later named *Trichomonasvirus* (TVV). This virus is believed to influence the host’s gene expression, impacting cytoadherence, drug susceptibility, and virulence proteins. However, studies on TVV-host interaction have produced conflicting results, with some genes appearing upregulated in one experiment but downregulated in another or vice versa. We hypothesize that these inconsistencies stem from the non-uniform distribution of TVV among individual cells within the same *T. vaginalis* isolate. Based on the experimental data from ultra-deep single-cell sequencing and immuno-localization, our study reveals that TVV’s intracellular distribution is highly heterogeneous and dynamic. These findings challenge the prevailing assumption that TVV is uniformly and statically distributed. This heterogeneity underscores the need to reassess previous research on TVV–*T. vaginalis* interactions, as earlier studies may have overlooked this crucial factor.

## Introduction

### Trichomonas vaginalis

*Trichomonas vaginalis* is an anaerobic, flagellated protozoan parasite that inhabits the human genitourinary tract and is the causative agent of trichomoniasis, the most common non-viral sexually transmitted infection (STI) globally. In 2016, the global prevalence of trichomoniasis was estimated at 5.3% among women and 0.6% among men. While the majority of infected individuals (70%–85%) are asymptomatic or experience only mild symptoms [1, 2], women are more likely to develop severe manifestations. These symptoms include inflammatory reactions, yellow vaginal discharge, vulvar irritation, and, in some cases, purulent vaginal discharge, vulvar or vaginal erythema, colpitis macularis, or even stillbirth [3]. Additionally, infection with *T. vaginalis* has been linked to a two-to threefold increased risk of HIV-1 transmission [4]. The primary treatment for trichomoniasis is metronidazole, a 5-nitroimidazole drug approved by the U.S. Food and Drug Administration (FDA) for oral or parenteral administration, achieving cure rates between 84% and 98% [5, 6]. However, an increasing number of clinical cases report *T. vaginalis* strains with reduced susceptibility to metronidazole [7–9], posing a significant threat to public health. This growing resistance highlights the urgent need to investigate factors contributing to reduced metronidazole susceptibility and to explore alternative treatment strategies to combat trichomoniasis effectively.

### Trichomonasvirus

*Trichomonasvirus* (TVV) is a double-stranded RNA (dsRNA) virus belonging to the Totiviridae family that infects *Trichomonas vaginalis* without causing lethal damage to its host [10]. The *Trichomonasvirus* genome is non-segmented, ranging from 4.5 to 5.5 kilobase pairs, and encodes two overlapping open reading frames for the capsid protein (CP) and RNA-dependent RNA polymerase (RdRp) [11]. Based on sequence variability, *Trichomonasvirus* is classified into five distinct species, designated as *Trichomonasvirus* 1 through 5 (TVV1-5) [10, 12]. Interestingly, it has been shown that a single *T. vaginalis* isolate can simultaneously harbor all five TVV species [12]. Despite the high sequence similarity of CP and RdRp across the species, the viral sequences confirm that these species are not products of mutations but distinct entities [12, 13]. The mechanism of TVV transmission remains unclear. Unlike many viruses, TVVs cannot transmit extracellular, as they lack the molecular machinery necessary for exiting and entering their protozoan host. The prevailing hypothesis suggests that TVVs are transmitted vertically during host replication via binary fission [10].

Various studies suggest that TVV infection significantly influences the gene expression profile of its host, *T. vaginalis*. The presence of the virus has been linked to increased adherence capacity, elevated levels of virulence-associated proteins, and enhanced drug resistance in infected parasites [14]. However, despite these observations, the clinical significance of TVV and its role in modulating the virulence and drug resistance of *T. vaginalis* remain poorly understood.

### Impact of *Trichomonasvirus* on the clinical symptoms of Trichomoniasis

TVV-infected parasites and purified *Trichomonasvirus* virions have been shown to enhance the dsRNA-dependent pro-inflammatory response. This response may exacerbate disease severity following metronidazole treatment, as the drug-induced damage to trichomonads can lead to a significant release of virions [15], potentially intensifying the host’s inflammatory reaction.

However, direct evidence connecting these in vitro findings to clinical outcomes is currently lacking. Table S1 summarizes the potential correlations between TVV infection and the clinical symptoms of trichomoniasis. However, the role of TVV in modulating the virulence and drug resistance of *T. vaginalis* remains poorly understood.

### Impact of *Trichomonasvirus* infection on host protein expression

Earlier research has shown that TVV infection increases the expression of cysteine proteinases [16] and the immunogen P270 [17] in *T. vaginalis*.

These cysteine proteinases enable the parasite to degrade components of the host vaginal epithelial cells, including hemoglobin, fibronectin, collagen IV, and the basement membrane. Additionally, cysteine proteinases play a vital role in *T. vaginalis* cytoadherence to the vaginal epithelium, contributing to both adherence [18–21] and cytotoxicity [22–24]. Secretory immunoglobulin A (SIgA), an antibody produced by cells in mucosal membranes, is a key component of host immunity. *T. vaginalis* infected with TVV has been found to upregulate the heterogeneous expression of Ig-degrading cysteine proteinases [25], enhancing the parasite’s cytoadherence, mediating cytotoxicity, and facilitating immune evasion. Given that protein regulation significantly influences host-pathogen interactions and pathogenicity, a deeper understanding of TVV-host interactions at the protein level is essential. Table S2 summarizes prior studies examining the effects of TVV infection on host protein expression

### Impact of *Trichomonasvirus* on Metronidazole Susceptibility

Metronidazole and tinidazole are currently the drugs of choice for treating trichomoniasis [25]. However, drug resistance to metronidazole has been reported in *T. vaginalis*. A previous study identified a correlation between a point mutation in the *T. vaginalis* genome and metronidazole resistance [26]. This mutation occurs at the 66th nucleotide of the internal transcribed spacer region, where thymidine replaces cytosine. Interestingly, this mutation is absent in trichomonads infected with TVV, suggesting that virus-infected parasites may exhibit increased susceptibility to metronidazole [26]. Despite this observation, the precise mechanism of how TVV influences metronidazole susceptibility remains unclear. Additionally, some studies have found no significant correlation between TVV and metronidazole resistance. These conflicting findings make it uncertain whether the virus plays a definitive role in modulating drug resistance. Table S3 summarizes previous research on the effects of *Trichomonasvirus* on metronidazole susceptibility.

The observed discrepancies in the impact of TVV on *T. vaginalis* biology highlight the need for a deeper understanding of the intricate interplay between TVV and *T. vaginalis*. Further research is crucial to unravel the mechanisms and implications of viral infection in *T. vaginalis*. Reaching definitive conclusions about virus-host interactions remains a significant challenge. One possible explanation for these conflicting findings lies in the heterogeneous distribution of TVV within *T. vaginalis* populations. Some Trichomonad cells within a single TVV-positive isolate may be TVV-free or infected with different TVV species, and this variability could account for the inconsistencies reported in previous studies. To investigate this hypothesis, we used agar colony culture to isolate individual *T. vaginalis* cells and applied ultra-deep single-cell RNA sequencing techniques to examine the distribution of TVV species across multiple *T. vaginalis* isolates. For the first time, our experimental results reveal that the distribution of TVV within the cells of a single *T. vaginalis* isolate is not homogeneous.

## Methods

### Trichomonas vaginalis culture

The *T. vaginalis* isolates (ATCC 30236, ATCC 50143, ATCC 50148, ATCC PRA-98) used in this study were obtained from the American Type Culture Collection (ATCC) in Manassas, Virginia, USA. The cells were initially cultured at a density of approximately 5 × 10⁵ cells/ml and maintained axenically in yeast extract, iron-serum (YI-S) medium at 37°C. For RNA-sequencing experiments, cells were harvested during the mid-exponential growth phase (1.8–2.3 × 10⁶ cells/ml). The viability of the cells was assessed using the trypan blue exclusion assay RNA Extraction and Quality Assessment

Total RNA was extracted from mid-log phase cultures of each *T. vaginalis* isolate using the GENEzol™ TriRNA Pure Kit (Geneaid Biotech Ltd., New Taipei City, Taiwan) following the manufacturer’s instructions. Briefly, cells were harvested from logarithmic phase cultures, the supernatant was discarded, and the cell pellet was washed with PBS. The pellet was then incubated with GENEzol reagent for 5 minutes. An equal volume of absolute ethanol was subsequently added and gently mixed. The mixture was transferred to RB Columns and centrifuged at 14,000 × g for 1 minute. The RB Columns were then sequentially rinsed with a pre-wash buffer followed by a wash buffer. Finally, purified RNA was eluted from the RB Columns using 20 µL of RNase-free water. RNA concentration was measured using a NanoDrop spectrophotometer (Thermo Fisher Scientific, Massachusetts, USA). The 230/280 and 260/280 absorbance ratios, calculated by the NanoDrop software, were used to assess RNA quality and potential protein contamination.

### Reverse Transcription-Polymerase Chain Reaction (RT-PCR)

The total RNA was reverse transcribed using the SuperScript™ III Reverse Transcriptase (Thermo Fisher Scientific, Massachusetts, USA). The target genes of *Trichomonasvirus* were amplified using species-specific primers: TVV1F2875-TVV1R3443, TVV2F2461-TVV2R3245, TVV3F61-TVV3R482, and TVV4F1338-TVV4R1834, as designed by Goodman et al. [27]. The amplification was performed with the 2× Super Hi-Fi Taq PCR MasterMix with loading dye (BIOTOOLS Co., Ltd., New Taipei City, Taiwan). Each reaction mixture (25 µL) contained 10 µM of each primer, 12.5 mL of the 2 × Super Hi-Fi Taq PCR MasterMix with loading dye, and 1 µg of the template derived from reverse transcription, following the manufacturer’s protocol. The thermal cycling conditions were as follows: inactivate reverse transcriptase at 94 °C for 3 minutes, followed by 30 cycles of 94 °C for 30 seconds, 55 °C for 30 seconds, and 72 °C for 1 minute, concluding with a final extension step at 72 °C for 5 minutes. The resulting amplified cDNA fragments were visualized after separation in a 1% agarose gel prepared with Tris-Acetate-EDTA (TAE) buffer with 0.01% FluoroVue™ Nucleic Acid Safe stain (SMOBIO Technology Inc., Hsinchu City, Taiwan).

### NGS RNA sequencing

The extracted RNA was used for library construction following a directional library preparation protocol. cDNA libraries were manually prepared using the Universal Plus™ mRNA-Seq with NuQuant® Kit (Tecan, Männedorf, Zürich, Switzerland) and sequenced on the Illumina system (Illumina, Inc., California, USA).

### Detection of Trichomonasvirus in Trichomonasvirus RNA-seq data

RNA-seq reads were aligned to the reference genomes of TVV species (TVV1-5) downloaded from the National Center for Biotechnology Information (NCBI) virus database. The analysis was performed using the RNA-Seq Analysis module in the CLC Genomics Workbench. Alignment parameters were set as follows: masking mode = no masking, match score = 1, mismatch cost = 3, linear gap cost model with insertion and deletion costs = 3, length fraction = 0.9, and similarity fraction = 0.8. The results section details the TVV species successfully aligned to their corresponding reference sequences among the *T. vaginalis* isolates.

### Indirect Immunofluorescent Assay

A total of 3 × 10˄6 trichomonads were inoculated onto a 0.1% Poly-L-lysine (Merck, Darmstadt, Germany) coated slide and incubated for 15 minutes at 37°C in an anaerobic chamber. After incubation, the medium was carefully removed, and the cells were fixed with 4% formaldehyde (FA) for 10 minutes at room temperature. The cells were then gently washed twice with 0.1% PBS-Tween20 (PBST) and permeabilized with 0.1% Triton X-100 in PBS, followed by blocking with 3% bovine serum albumin (BSA). Next, the cells were incubated for 1 hour at room temperature with the anti-dsRNA [9D5] primary monoclonal antibody (Absolute Antibody Ltd, Cleveland, UK) at a 1:1200 dilution in blocking solution. Following two washes with PBST, a goat anti-rabbit secondary antibody conjugated with Alexa Fluor 594 (Thermo Fisher Scientific Inc., Massachusetts, USA) was applied at a 1:1000 dilution, along with DAPI (4’,6-diamidino-2-phenylindole). The sample was incubated for an additional hour at room temperature. After two more washes with PBST, the slides were air-dried, and 4 μL of FluoroQuest™ Antifade Mounting Medium (AAT Bioquest, Inc., California, USA) was applied to mount the slides. Stained slides were visualized using a ZEISS LSM510META NLO Microscope (ZEISS, Stuttgart, Germany).

### Single-cell RNA-seq and data analysis

Single-cell RNA sequencing was performed on *T. vaginalis* cells harvested during the mid-log growth phase. Cell viability and concentration were assessed using trypan blue staining. Following 10× Genomics’ guidelines for targeting 1,000 cells, the appropriate cell volume was loaded onto a 10× Genomics Chromium GEM Chip (10x Genomics Inc., Pleasanton, CA, USA) and processed in the Chromium Controller for cell capture and library preparation. Microfluidics technology was used to achieve cell capture and library preparation by combining cells with Single Cell 3’ Gel Beads containing barcoded primers with unique molecular identifiers (UMIs).

The workflow included cell lysis, barcoded reverse transcription of RNA, amplification of barcoded cDNA, fragmentation of cDNA into 200 bp fragments, addition of 5’ adapters, and sample indexing. All procedures followed the manufacturer’s protocol, using the Next GEM Single Cell 3’ Reagent Kits v3.1_LT (Dual Index). Prepared libraries were sequenced on the Illumina NovaSeq 6000 / Illumina NovaSeq X Plus platform (Illumina, Inc., San Diego, CA, USA).

Preliminary data analysis was conducted using the Cell Ranger pipeline (10x Genomics Inc., Pleasanton, CA, USA) on a Linux system. The reference library was created by combining whole-genome reference sequences and annotation files for TVV and *T. vaginalis* obtained from the NCBI FTP server. A manually curated annotation file for all chromosomes and contigs based on individual gene information was also included. Sequencing reads were aligned against the reference library, and the outcomes were further analyzed and visualized using R packages Seurat v5.1.0 [28], Monocle v1.3.7[29], and UpSetR [30]. Viral species infection categorization was primarily based on the detection of virus-specific glycoprotein gene signals.

## Results

### Characterization of *Trichomonasvirus* Infection in *Trichomonasvirus* Isolates

To confirm the presence of *Trichomonasvirus* (TVV) in various *T. vaginalis* isolates, RNA sequencing (RNA-seq) data were mapped to reference sequences of *Trichomonasvirus*. The infection status of 24 isolates was systematically analyzed and summarized in Table 1. Among these, the isolate ATCC 50143 was confirmed to be virus-free, whereas ATCC 30236 was infected by TVV1 only. Notably, double infections were detected in isolate ATCC PRA-98 (harboring TVV2 and TVV3), while a triple infection was observed in isolate ATCC 50148 (harboring TVV1, TVV2, and TVV3) (Fig. 1A–B). These four isolates were selected for further investigation. Initial validation via polymerase chain reaction (PCR) (Fig. 1C) confirmed findings consistent with the in silico RNA-seq analysis, reinforcing the reliability of the sequencing approach in viral identification.

**Fig 1.**
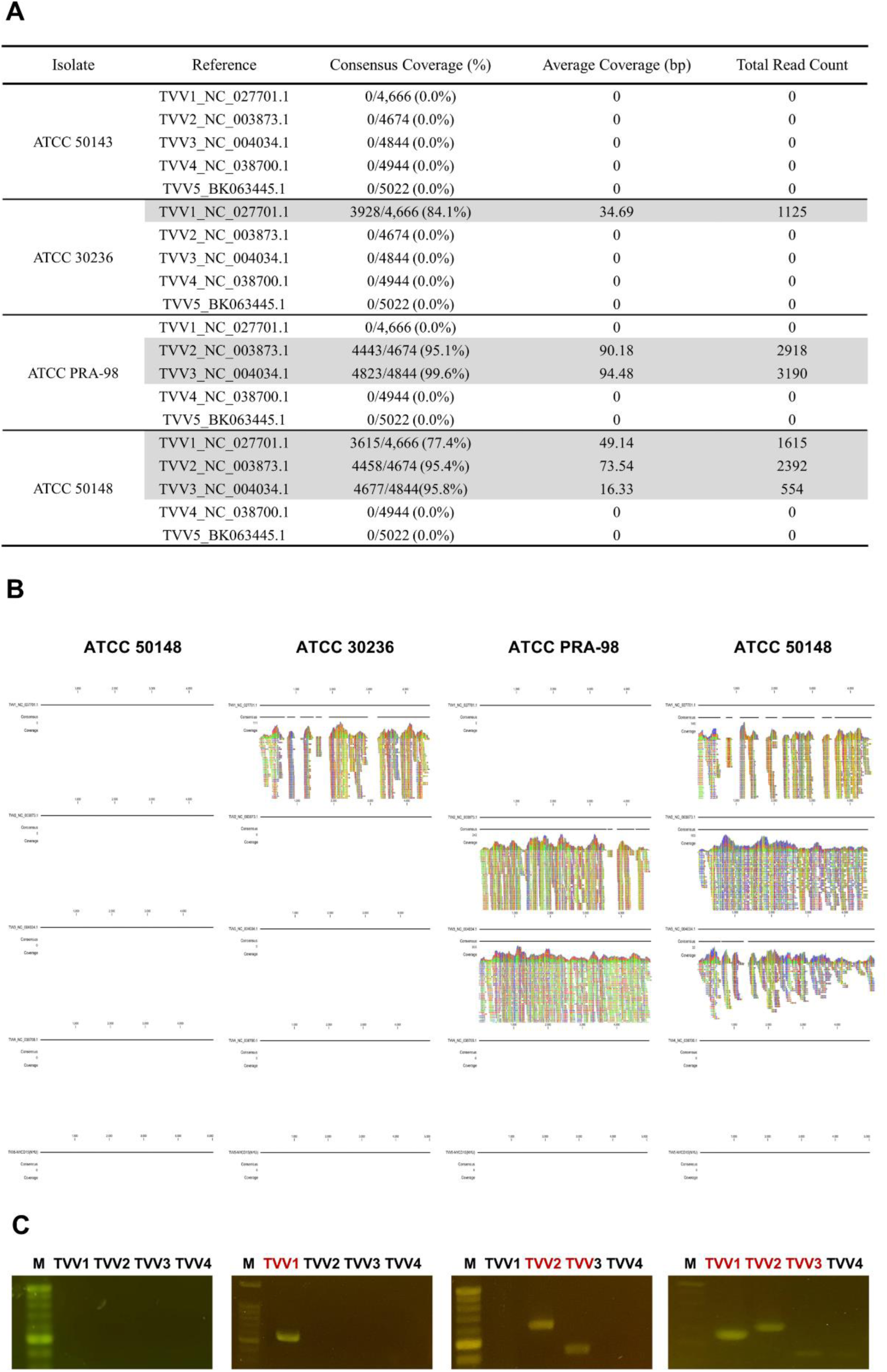
*Trichomonasvirus* species infection in *T. vaginalis* isolates. The presence of TVV in the *T. vaginalis* isolates (ATCC 30236, ATCC 50143, ATCC 50148, ATCC PRA-98) was determined by mapping RNA-seq reads to *Trichomonasvirus* reference sequences obtained from the NCBI Virus Database using QIAGEN CLC Genomics Workbench (v20.0.3). (A) Summary of consensus coverage, average coverage, and total read counts. (B) Schematic representation of the consensus contig alignment for each *Trichomonasvirus* species. (C) Detection of *Trichomonasvirus* isolates using RT-PCR with virus species-specific primers, with PCR products separated on a 1% agarose gel.

**Table 1.** *Trichomonasvirus* species infected in *Trichomonasvirus* isolates.

| Isolate | Number<br>of<br>Species | TVV1 | TVV2 | TVV3 | TVV4 | TVV5 |
| --- | --- | --- | --- | --- | --- | --- |
| NYCB20 | 0 |  |  |  |  |  |
| ATCC 50143 (CDC085) | 0 |  |  |  |  |  |
| ATCC 50167 (B7RC2) | 0 |  |  |  |  |  |
| T1 | 1 | ● |  |  |  |  |
| B7268 | 1 | ● |  |  |  |  |
| ATCC 30236 (JH 31A #4) | 1 | ● |  |  |  |  |
| T016 | 1 |  | ● |  |  |  |
| ATCC 30001 (C-1:NIH) | 1 |  | ● |  |  |  |
| ATCC 30238 (JH 32A #4) | 1 |  |  | ● |  |  |
| GOR69 | 2 | ● | ● |  |  |  |
| NYCF20 | 2 | ● |  | ● |  |  |
| ATCC 50142 (RU393) | 2 | ● |  | ● |  |  |
| LSU180 | 2 | ● |  |  | ● |  |
| CDC1103 | 2 | ● |  |  | ● |  |
| CDC1123 | 2 | ● |  |  | ● |  |
| CDC1132 | 2 | ● |  |  | ● |  |
| NYCG31 | 2 | ● |  |  |  | ● |
| PRA-98 (G3) | 2 |  | ● | ● |  |  |
| ATCC 50148 (NYH286) | 3 | ● | ● | ● |  |  |
| NYCA04 | 3 | ● |  | ● |  | ● |
| NYCE32_8 | 3 | ● |  | ● |  | ● |
| SD2 | 3 | ● |  | ● |  | ● |
| NYCC37 | 4 | ● | ● | ● |  | ● |
| NYCD15 | 5 | ● | ● | ● | ● | ● |

### Heterogeneous Distribution of *Trichomonasvirus* Within Single *T. vaginalis* Isolates

To investigate the distribution of *Trichomonasvirus* within *T. vaginalis* isolates, single-cell RNA sequencing (scRNA-seq) was employed. The sequencing data were processed using the Cell Ranger pipeline (10x Genomics Inc., Pleasanton, CA, USA), followed by downstream analysis using Seurat v5.1.0 and Monocle v1.3.7 in R. These tools enabled the classification of individual cells based on viral protein expression patterns. Visualization of the viral distribution in three virus-infected isolates revealed significant heterogeneity in the presence of virus among cells (Fig. 2A). This observation was further validated by immunofluorescence assays, which demonstrated specific binding of a dsRNA-specific antibody to viral-derived dsRNA within the cytoplasm (Fig. 2B). Collectively, these findings confirm that *Trichomonasvirus* exhibits an uneven distribution at the single-cell level within a given isolate.

**Figure 2.**
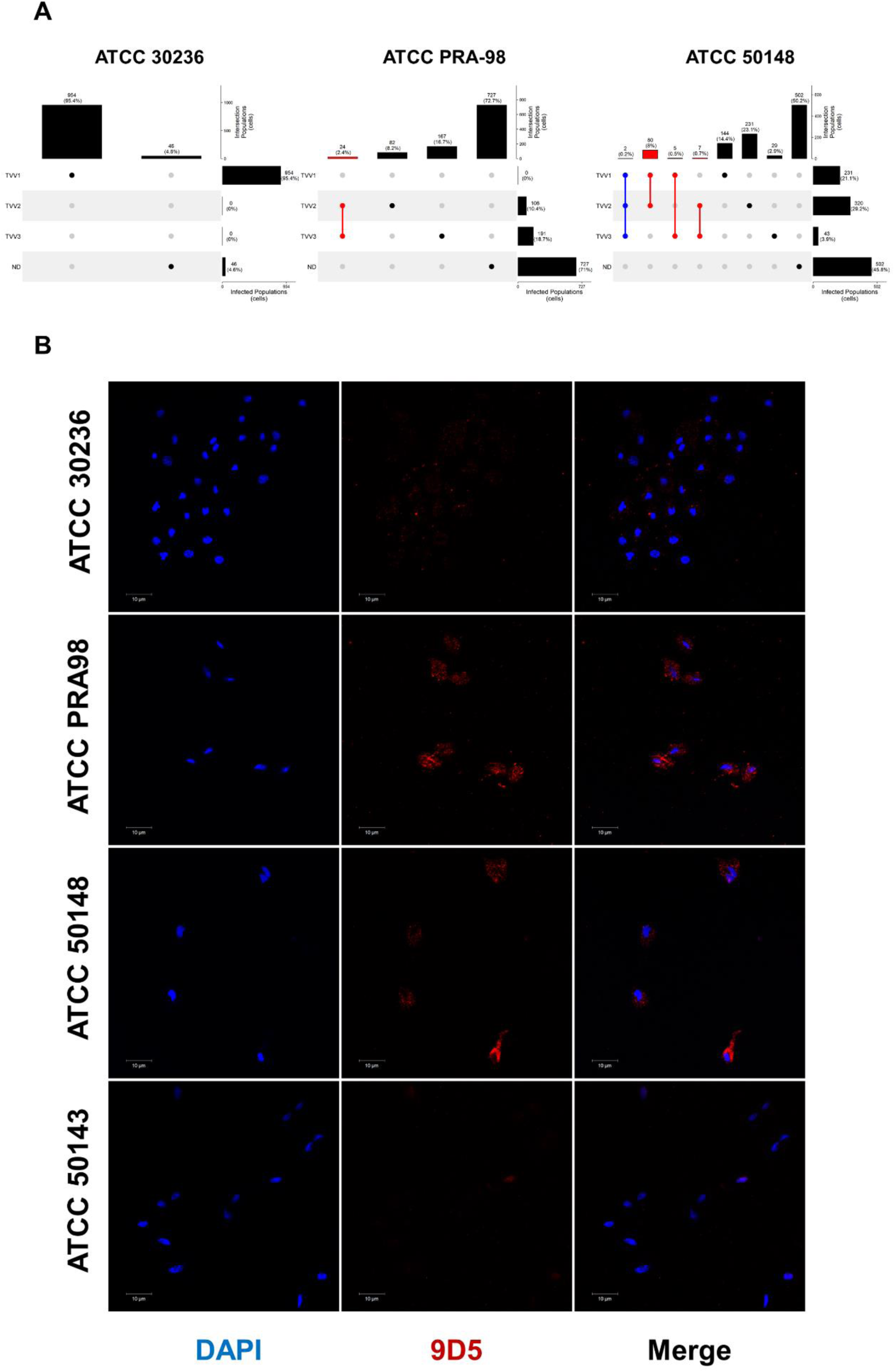
Heterogeneous distribution of *Trichomonasvirus* in different *T. vaginalis* isolates. (A) Upset plots illustrate the distribution of viral populations among the *T. vaginalis* isolates (ATCC 30236, ATCC PRA-98, ATCC 50148, and ATCC 50143) using scRNA-seq. The plots show the presence of *Trichomonasvirus* species and their co-occurrence patterns across isolates. The bar graphs indicate the total number of reads mapped to each viral species in individual isolates. (B) Detection of viral dsRNA using the dsRNA-specific antibody 9D5 in different *T. vaginalis* isolates. Nuclei were stained with DAPI (blue), while viral dsRNA was detected using the 9D5 antibody conjugated with Alexa Fluor 594 (red). Images indicate the heterogenicity of viral dsRNA within the parasite. Scale bars = 10 µm.

### Heterogeneity in Viral Load Across Individual Cells

To further characterize the heterogeneity of viral infection, we analyzed both the presence of viral transcripts and the variation in viral load among individual cells. The ATCC 30236 isolate, which harbors a single TVV1 infection, was subjected to a higher sequencing depth to assess viral expression dynamics. The results demonstrated substantial variability in the expression levels of viral-associated proteins among virus-positive cells, with some exhibiting significantly higher viral loads than others (Fig. 3). These findings highlight the heterogeneous nature of viral replication and maintenance within single-cell populations.

**Fig 3.**
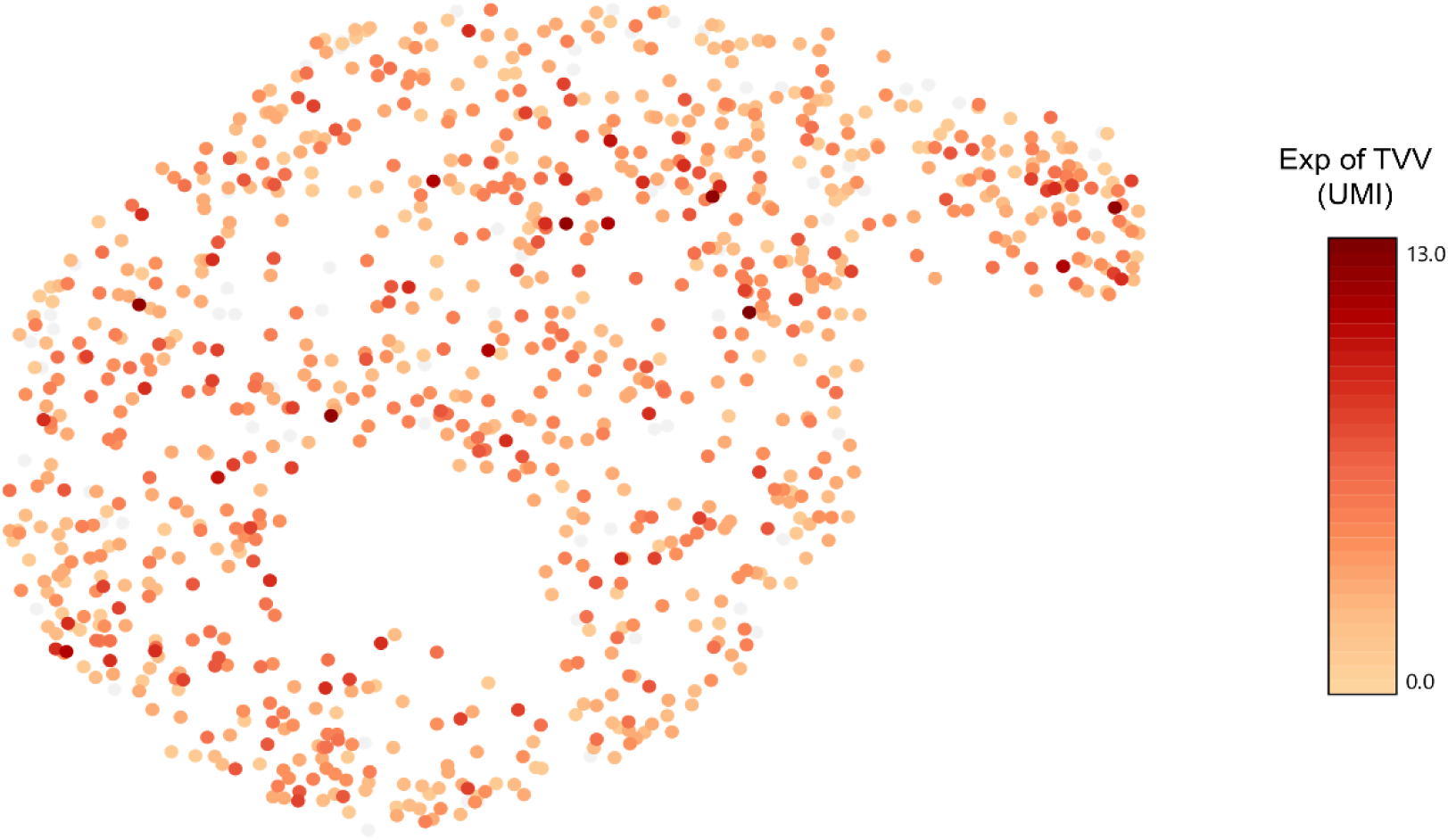
Heterogeneity Expression of Viral Protein in Single *T. vaginalis* Isolate. High-resolution scRNA-seq data from the *T. vaginalis* isolate ATCC 30236 was visualized using Loupe Browser 8.0.0, with uniform manifold approximation and projection (UMAP) dimensionality reduction. Each point represents an individual cell, with color intensity indicating the expression levels of viral-associated proteins, quantified as unique molecular identifiers (UMI). The scale bar reflects the transcript abundance distribution, highlighting the heterogeneity in viral gene expression across the population.

### Dynamic Temporal Changes in Viral Distribution Within Host Cells

To determine whether viral distribution remains static within a given isolate over time, the ATCC PRA-98 isolate, which harbors double infections (TVV2 and TVV3), was subjected to two additional independent scRNA-seq experiments at different time points, facilitating a triplet comparative analysis. A chi-square test was performed to assess statistical differences between these datasets. The results revealed significant temporal variation in viral distribution across single-cell populations (Fig. 4), suggesting that *Trichomonasvirus* infection does not remain constant but instead undergoes dynamic changes within host cells over time. This temporal heterogeneity underscores the complex interplay between viral persistence, host cellular mechanisms, and potential viral transmission dynamics.

**Fig 4.**
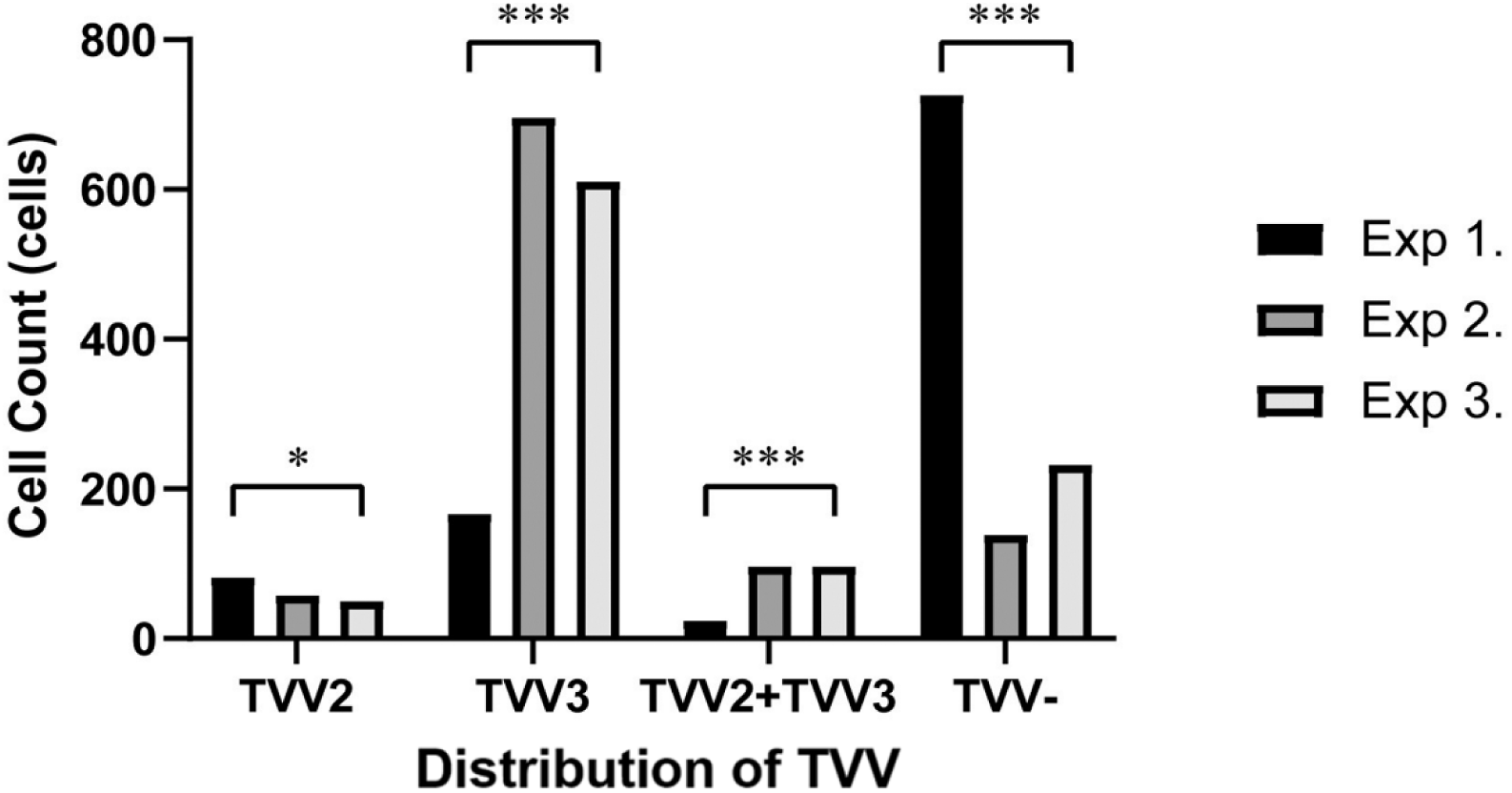
Temporal Dynamics of *Trichomonasvirus* (TVV) The distribution of *Trichomonasvirus* (TVV) over time was analyzed and quantified using the Seurat v5.1.0 R package. Cell counts corresponding to the distribution of TVV were compared across three independent experiments (Exp. 1, Exp. 2, and Exp. 3). A chi-square test was conducted to evaluate statistically significant differences among groups, with significance levels denoted (*p < 0.05, ***p < 0.001).

## Discussion

Our study provides compelling evidence that the distribution of TVV within *T. vaginalis* is inherently heterogeneous and dynamic, challenging the long-held assumption of uniform viral distribution. This discovery highlights the need to revisit established paradigms regarding TVV-host interactions and calls for deeper investigations into the mechanisms underlying this variability.

Through single-cell sequencing of TVV-positive *T. vaginalis* isolates, we revealed that a significant proportion of cells remain uninfected, even within the same isolate. This heterogeneity, or the ratio of infected/uninfected cells, points to a complex and nuanced relationship between the virus and its host, raising important questions about the factors governing infection dynamics. Notably, our findings address previous concerns about the potential loss of *Trichomonasvirus* during prolonged laboratory cultivation [32]. Contrary to these concerns, we observed no evidence of viral disappearance under such conditions. However, even within TVV-positive isolates, many cells do not contain any virus, suggesting that viral transmission is a dynamic process. A plausible explanation for this phenomenon, supported by prior research, is the packaging of *Trichomonasvirus* particles into extracellular vesicles, enabling their release and exchange among host cells [33, 34].

Our findings also go beyond the binary presence or absence of viral infection, uncovering significant variability in viral load among infected cells. While some cells exhibited high levels of viral RNA, others displayed markedly lower amounts, even within the same isolate. This uneven distribution suggests that cell-specific factors, such as the cellular environment or infection cycle stage, influence viral replication and persistence. These results underscore the importance of single-cell resolution analyses to fully capture the complexity of *Trichomonasvirus*-host interactions.

Additionally, our results provide context for earlier studies demonstrating the significant impact of *Trichomonasvirus* on clinical symptoms and the transcriptomic and proteomic profiles of *T. vaginalis*. These studies often relied on the assumption of uniform viral distribution—a premise that our findings suggest requires reevaluation. Incorporating viral heterogeneity into experimental designs is essential for accurately interpreting the interplay between the virus and its host and understanding this variability’s broader implications. Moreover, the conflicting results regarding the viral impact (Tables S1–S3) may be attributed to differences in viral load or isolate-specific properties. To address this, our group is establishing a novel system for studying the impact of *Trichomonasvirus*, wherein a 2CMC-treated virus-positive isolate [41] is reinfected using extracellular vesicles [33, 34]. This approach allows us to eliminate the influence of isolate-specific properties.

Furthermore, adjusting the concentration of extracellular vesicles during reinfection can precisely control the viral load before conducting viral impact assays, enabling a more targeted investigation of TVV’s effects on its host.

Our study demonstrates the highly dynamic nature of *Trichomonasvirus* (TVV) distribution, highlighting the necessity of confirming the proportion of TVV-positive cells and even quantifying the viral load within individual cells before conducting experiments. The immunofluorescence assay (IFA) serves as a powerful tool for this purpose, allowing us to identify virus-positive populations by counting fluorescence-positive cells. Additionally, fluorescence signal intensity can provide an estimate of viral load. However, in the virus-negative isolate ATCC 50143 (Fig. 2B), a low-level dsRNA signal was detected, likely originating from short endogenous dsRNA motifs in tRNA-derived small RNAs (tsRNAs) or endogenous small interfering RNAs (endo-siRNAs) [35, 36]. This finding underscores the need for more stringent criteria to distinguish between truly virus-negative populations and those with low viral loads.

Nevertheless, by recognizing the dynamic and heterogeneous nature of *Trichomonasvirus* distribution, our study lays the groundwork for future research to elucidate these interactions’ biological and pathogenic significance. These insights offer an opportunity to refine our understanding of *Trichomonasvirus* biology and its role in the pathogenicity of *T. vaginalis*, paving the way for innovative approaches to studying this complex system.

## Acknowledgments

We would like to thank the Microscopy Center at Chang Gung University for its technical assistance in laser confocal microscopy and Min-Chi Chen from the Department of Public Health and Biostatistics Consulting Center, School of Medicine, Chang Gung University, for assistance of biostatistical advisement. This research was supported by Chang Gung Memorial Hospital (CMRPD1J0311∼1J0CMRPD1J0311∼1J0313313, CMRPD1M0571∼1M0572) and National Science and Technology Council (NSTC 113-2320-B-182-017-MY3). H.W.L received support from Ministry of Education (MOE) Doctoral Scholarships.

## Data Availability

Raw sequencing data are available in the Sequence Read Archive (SRA). The scRNA-seq data have been submitted under BioProject PRJNA1238186, bulk RNA-seq data from our lab have been submitted under BioProject PRJNA1241774. Other publicly available datasets were retrieved from the following accessions: NYCB20A, B7268, ATCC 30236, GOR69, NYCF20, NYCG31, NYCA04, SD2, NYCC37, NYCD15 (PRJNA280779), T016 (SRR585698), and NYCE32_8 (SRX1686500).

## Supplementary data

**Table S1.** TVV Effects on *Trichomonasvirus* Proteins Expression.

| **isolate (+/-/TVV subspecies)** | **Descriptions** |
| --- | --- |
| TVV(+): JHHR, ATCC30001, JH31A<br>TVV(-): CDC85, IR78, RU375,<br>DD3Cz, 5147Cz, AL8WCz, AL11Cz,<br>355Cz, 427Cz, JWCz | TVV+ is linked to the ability of<br>*Trichomonasvirus* to display<br>immunogens on their surfaces and<br>undergo phenotypic variation. [37] |
| TVV(+): AL8, AL10, 347, 8111 | P270 mRNA accumulates exclusively in<br>TVV+ trichomonads, resulting in<br>transcriptional expression.<br>Consequently, low P270 protein levels<br>are present in TVV- progeny. [17] |
| TVV(+): TO56<br>TVV(-): T068-II | TVV can enhance immunoglobulin<br>degrading activity by expressing viral<br>products, including potential proteinase<br>or upregulating trichomonad proteinase.<br>[25] |
| TVV(+): 21201, T068-II, NYH286,<br>AL8, AL10, 347, 8111 | Significant variations in protein and<br>proteinase profiles between TVV+<br>parent and TVV- progeny organisms.<br>[38] |
| - | TVV+ and TVV- isolates showed 50<br>differentially expressed proteins. [39] |

**Table S2.**
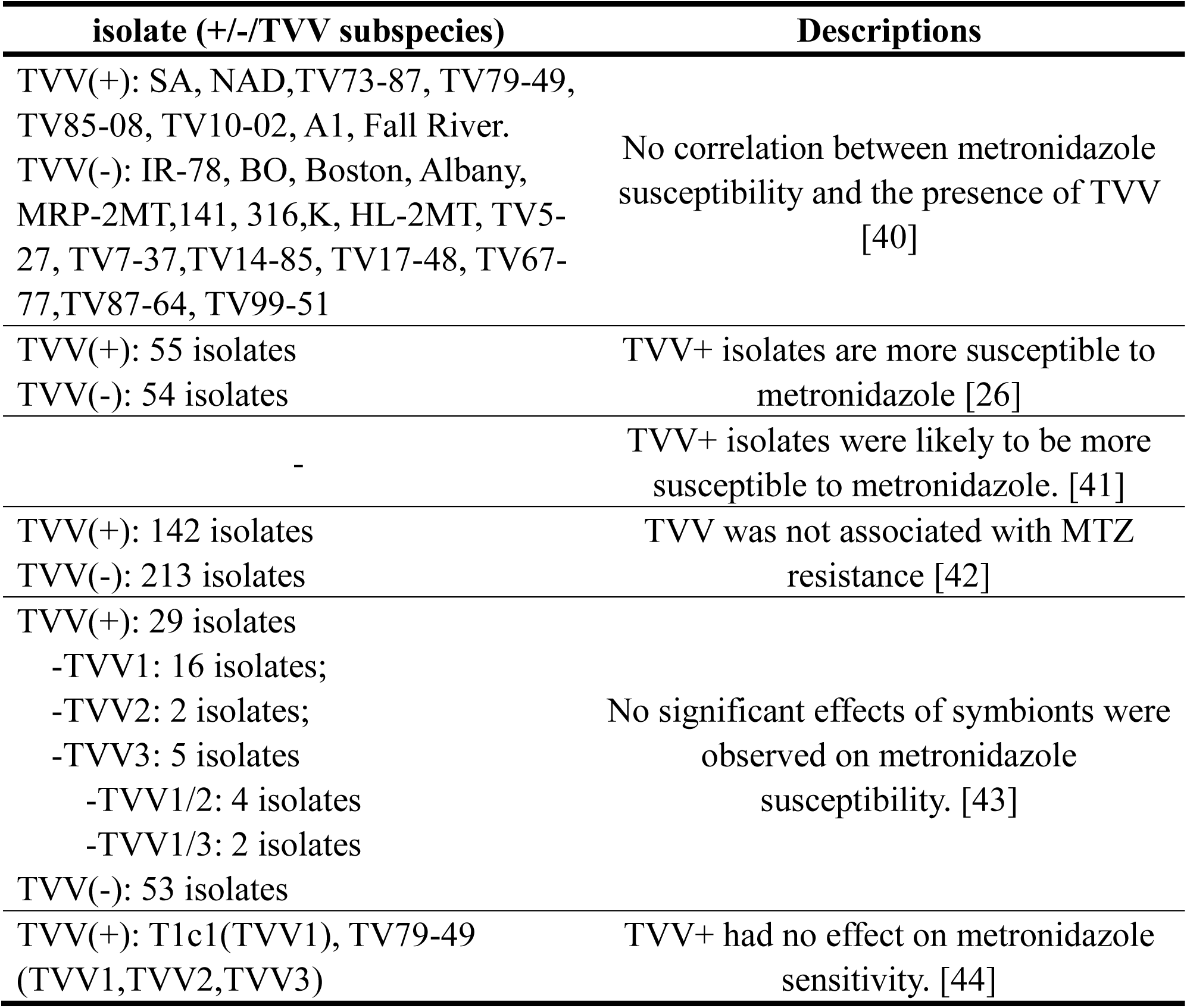
TVV Effects on *Trichomonasvirus* Metronidazole Susceptibility.

**Table S3.** TVV Effects on *Trichomonasvirus* Clinical Symptoms.

| isolate (+/-/TVV subspecies) | Descriptions |
| --- | --- |
| TVV(+): NYH286,302, 287, 370, 366, 323, 232, 335<br>TVV(-): T016, 433 | TVV+ isolate is associated with higher parasite virulence in the vaginal microenvironment. [45] |
| TVV(+): IH-1, IH-2, IH-3<br>TVV(-): IH-6, IH-8, IH-10, IH-11 | Isolates with weak pathogenicity in mice linked to persistent TVV presence. [46] |
| TVV(+): 22 isolates<br>TVV(-): 18 isolates | TVV+ were associated with more frequent vaginal discharge, dysuria, and cervical erythema, dyspareunia. [47] |
| - | TVV + caused differences in symptomatology, drug resistance, parasite load, and transmissibility. [10] |
| TVV(+): UR1(TVV1) | TVV altered epithelial responses linked to the type of vaginal bacteria present during trichomoniasis. [48] |
| TVV(+): A170(TVV1), C334(TVV1), C344(TVV1), A5(TVV1/2), A42(TVV2), A69(TVV1), C76(TVV2), A185(TVV1), C9(TVV1), C206(TVV1,TVV2), C240(TVV2), C247(TVV1/2), C187(TVV2), A66(TVV2), C173(TVV2), C308(TVV2), C313(TVV2), C349(TVV2), C351(TVV2), C352(TVV2), C355(TVV2)<br>TVV(-): C91(-), C129(-), C175(-), C237(-), C239(-), C350(-), C350(-), C356(-), C358(-), C361(-), A59(-), C15(-), C98(-), C190(-),A47(-),C348(-), | TVV2 was associated with higher cytoadhesion. [49] |
| TVV(+): 142 isolates<br>TVV(-): 213 isolates | TVV was not associated with clinical symptoms, repeat TV infections. [42] |
| TVV(+): 29 isolates<br>-TVV1: 16 isolates;<br>-TVV2: 2 isolates;<br>-TVV3: 5 isolates<br>-TVV1/2: 4 isolates<br>-TVV1/3: 2 isolates<br>TVV(-): 53 isolates | No significant effects of symbionts were observed on cytotoxicity or symptom severity. [43] |

## Notes

### Competing Interest Statement

The authors have declared no competing interest.

